# Restricting a single amino acid cross-protects *D. melanogaster* from nicotine poisoning through mTORC1 and GCN2 signalling

**DOI:** 10.1101/2021.11.03.467051

**Authors:** Tahlia L. Fulton, Christen K. Mirth, Matthew D. W. Piper

## Abstract

Dietary interventions that restrict protein intake have repeatedly been shown to offer beneficial health outcomes to the consumer. Benefits such as increased stress tolerance can be observed in response to restricting individual amino acids, thus mimicking dietary protein restriction. Here, we sought to further understand the relationship between dietary amino acids and stress tolerance using *Drosophila melanogaster*. Utilising a chemically defined medium for *Drosophila*, we found that transiently restricting adult flies of a single essential amino acid generally protects against a lethal dose of the naturally occurring insecticide, nicotine. This protection was conferred during the pre-treatment window, was specific for individual amino acids and depended on the identity of the focal amino acid, as well as the duration and intensity of its restriction. For instance, complete isoleucine deprivation for 7 days maximised its protective effect - increasing survival during nicotine exposure by 100%. However, a dose of 25% threonine was required to maximise its protective effect (53% enhanced survival). To understand the molecular basis of these effects, we modified the signalling of two cellular sensors of amino acids, GCN2 (General control non-derepressible) and mTORC1 (mechanistic Target of Rapamycin Complex 1) in combination with amino acid restriction. We found that GCN2 was necessary for diets to protect against nicotine, whereas suppression of mTORC1 was sufficient to induce nicotine resistance. This finding implies that amino acid restriction acts via amino acid signalling to cross-protect against seemingly unrelated stressors. Altogether, our study offers new insights into the physiological responses to restriction of individual amino acids that confer stress tolerance. This has broad potential for application in animal and human health.

## Introduction

Nutrition is a powerful regulator of health, and manipulating the quantity and quality of diet affects fitness traits such as longevity, reproduction, and stress resistance (1-3). Transient dietary restriction, which is defined as reduced nutrient intake without malnutrition, increases stress resistance, improves metabolic health, and extends lifespan across a broad range of organisms (1, 4-6). Furthermore, restricting dietary protein alone is sufficient to mimic the benefits of restricting food intake (5, 7-9). These effects can also be mimicked by restricting or removing single dietary amino acids. For instance, methionine restriction enhances tolerance to chemical and thermal stress in yeast, mice, and human cells (10-12), and transient deprivation of tryptophan has also conferred protection against a model of surgical stress in mice (13). These effects of diet have attracted interest for their potential to enhance lifelong human health. Current data indicates an important role for the evolutionarily conserved amino acid sensing pathways in mediating these effects (7, 13), though little is known about the downstream steps that are required to confer protection or the breadth of stress resistance they afford.

The presence or absence of dietary amino acids is detected and signalled by the complementary effects of the intracellular kinases, GCN2 (General Control Non-derepressible 2) and mTORC1 (mechanistic Target Of Rapamycin Complex 1), which are highly conserved and well characterised for their role in growth, metabolism, and lifespan (8). In the presence of amino acids GCN2 is inactive, but mTORC1 is activated and signals to a cascade of downstream effectors that promote protein translation and growth (14). In contrast, when amino acids are absent, GCN2 is activated and mTORC1 is suppressed, ultimately resulting in reduced global translation and the promotion of autophagy; a process whereby cellular components are engulfed and broken down to be recycled for use in the synthesis of selected proteins important for stress resistance (15, 16). As amino acids are recycled via autophagy, their presence has the potential to reactivate mTORC1 despite the ongoing nutrient shortage. To ensure that mTORC1 remains inactive during nutrient scarcity, GCN2 also sustains mTORC1 suppression during prolonged amino acid deprivation (17). These responses to amino acid shortage are important for prioritising survival over growth when nutrients are scarce.

The full extent of these signalling interactions and the specific molecules by which each of the individual amino acids are detected are still being uncovered. Some essential amino acids, such as methionine, leucine, and arginine, have been directly linked to molecular sensors upstream of mTORC1 (18-20). However, it hasn’t yet been determined whether there are other amino acids that regulate mTORC1 activity *in vivo*. In contrast, GCN2 is thought to detect intracellular depletion of any amino acid by binding uncharged tRNAs, which accumulate when there are insufficient free cognate amino acids (21). GCN2 can also suppress mTORC1 activity through multiple effectors (17, 22), meaning that ultimately, the balance of activity of both of these kinases is sensitive to the levels of any single amino acid.

As the mechanisms for amino acid detection are highly conserved across eukaryotes. This makes animals like the fruit fly *Drosophila melanogaster*, with their short generation times and abundance of genetic tools, excellent models to understand the relationship between dietary amino acids, amino acid signalling, and their roles in stress tolerance. Particularly for fruit flies, genetic and pharmacological tools can be coupled with a chemically defined (holidic) medium to investigate how restricting individual dietary amino acids confers tolerance to stressors (23).

A group of ecologically relevant stressors for flies are toxic plant-derived insecticides. These molecules are metabolised and excreted by the drug detoxification system, which consists of three evolutionarily conserved phases (24). Phase I and II metabolism catalyse modifications of toxins into more hydrophilic molecules that are more easily excreted in phase III metabolism (25). The superfamilies of genes involved in these phases of detoxification include CYP450 (Cytochrome P450) monooxygenases, GSTs (Glutathione S-transferases), UGTs (UDP-Glucuronyl transferases), and ABC (ATP-binding cassette) transporters (25, 26). Since xenobiotic toxins are ingested as part of regular food consumption, it is reasonable to anticipate that there is a link between nutrition and the need to detoxify these compounds. We therefore hypothesised that food composition, in particular amino acid restriction, could be an effective tool to modify resistance to chemical stressors and that this may be mediated by evolutionarily conserved nutrient sensors. If so, understanding how dietary interventions protect organisms against endobiotic and xenobiotic toxins would be useful in understanding the role of diet in animal, and therefore human, health.

Here we found that transient deprivation of almost any essential amino acid increased nicotine resistance in *Drosophila*. However, the magnitude of this dietary protection depended on the duration of exposure to the amino acid depleted diet, the identity of the amino acid being manipulated, and to what extent the amino acid was being restricted. We also found that this protection was driven by interactions between GCN2 and mTORC1. These data highlight a new role for dietary amino acids in modifying the stress tolerance of *Drosophila* and reveals new insights into how this protection is effected.

## Results

### Removing an essential amino acid from the diet can protect flies against future toxin stress

In previous studies, restriction of dietary protein or individual amino acids has conferred stress tolerance in mice (5, 7, 13). To identify the mechanisms through which amino acids confer protection, we first sought to examine if this phenomenon also occurred in flies. To do this, we tested if short-term exposure to individual amino acid dropout diets could protect flies against exposure to nicotine, a chemical stressor that is detoxified by the evolutionarily conserved drug detoxification system (27) (Figure 1.A).

**Figure 1.**
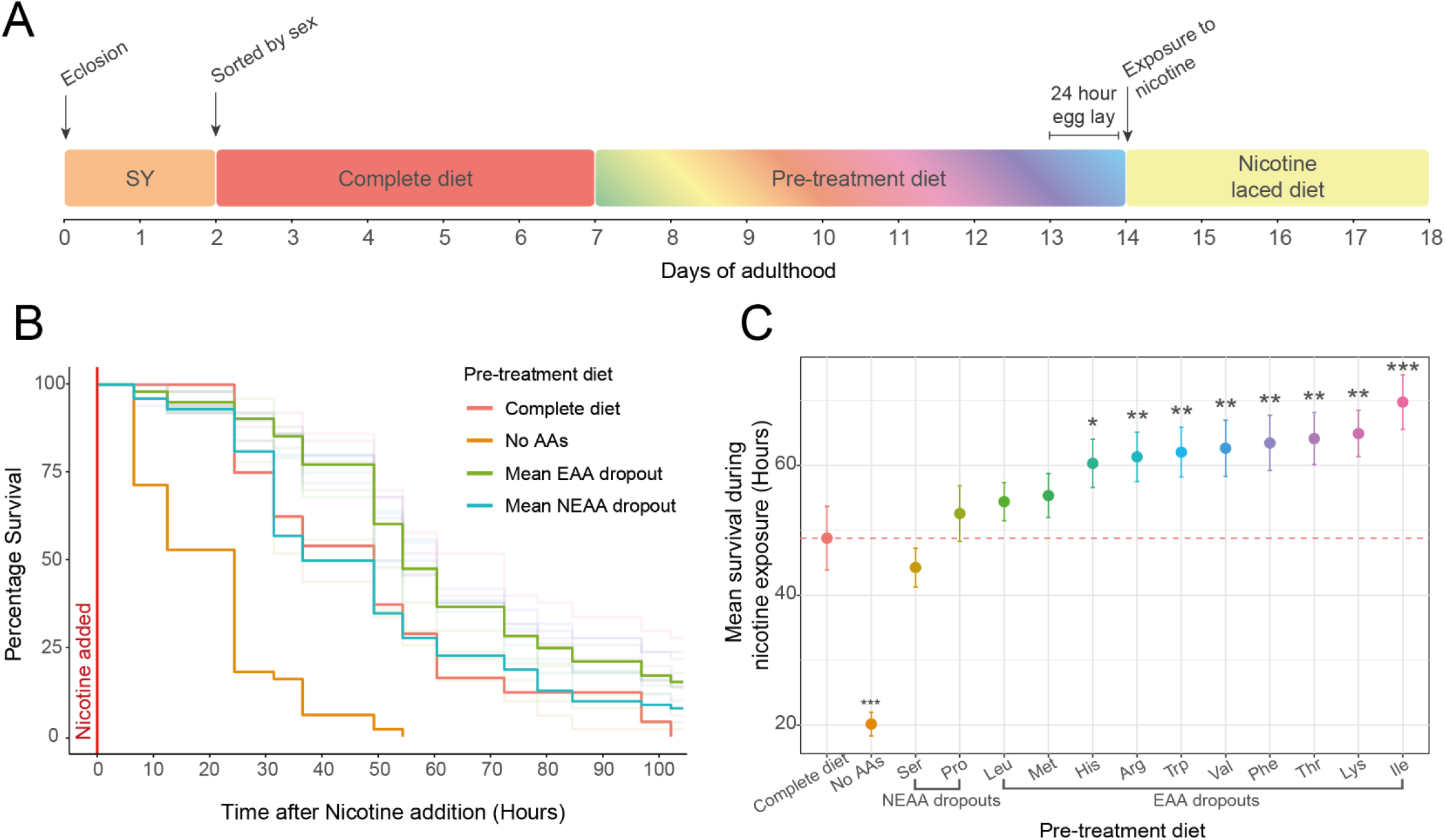
Removing an essential amino acid from the diet can protect flies against future toxin stress. (**A**) Flies were pre-treated with one of 14 diets for 7 days before chronic exposure to food containing 1mg/mL nicotine. Diets included a complete diet, a diet lacking all amino acids, a serine or proline dropout and dropouts for each essential amino acid (**B**) Survival curves of pre-treated flies once exposed to nicotine; bold lines represent the average survival of the transparent lines, which represent individual dropout diets. (**C**) Mean survival +/- SEM of pre-treatment groups; groups were compared to the complete diet (red). N = 50 flies that were pre-treated with a complete diet and N = 100 for all other pre-treatment groups.

Firstly, to assess the effect of our pre-treatment diets on fly physiology, we measured the flies’ fecundity immediately prior to nicotine exposure. Flies fed a complete diet or either one of two non-essential amino acid dropouts had equally high levels of egg laying (P > 0.98; Figure S1.A), whereas flies fed any one of the ten essential amino acid dropouts displayed severely reduced fecundity (P<0.001) that was no different from flies fed a diet that lacked all amino acids (P > 0.5). These data confirm previous observations on the requirement for these amino acids for egg production (23, 28)

Following 7 days of pre-treatment on our dropout diets, flies were exposed to food containing 1mg/mL nicotine and their survival was recorded. This dose of nicotine caused most flies to die within 5 days, whereas flies that were maintained without nicotine exhibited no mortality during this time (Figure S1.B), indicating that the pre-treatment diets were not causing death. Interestingly, we found that eight of the ten essential amino acid dropout diets conferred protection against nicotine treatment when compared to the complete diet control (P < 0.02, Figure 1.B & C). By contrast, flies fed either of the non-essential amino acid dropout diets or a diet lacking either of the essential amino acids leucine or methionine were no different from those on a complete diet (P > 0.1). Finally, flies pre-treated with food lacking all amino acids had worse survival upon nicotine exposure than complete diet controls (P < 0.001; Figure 1.B & C). These results indicate that some single amino acid dropouts can protect flies against subsequent toxin stress and that the degree of protection is specific to the amino acid dropout they experience.

### Pre-treatment modifies responses to high doses of nicotine

To understand the range of nicotine protection conferred by amino acid dropout diets, we pre-treated flies with either a complete diet, an isoleucine dropout, or a diet lacking all amino acids, and subsequently exposed them to nicotine concentrations between 0 and 1mg/mL (Figure 2A). We chose the isoleucine dropout diet for this experiment as it previously showed the greatest protection against nicotine.

**Figure 2.**
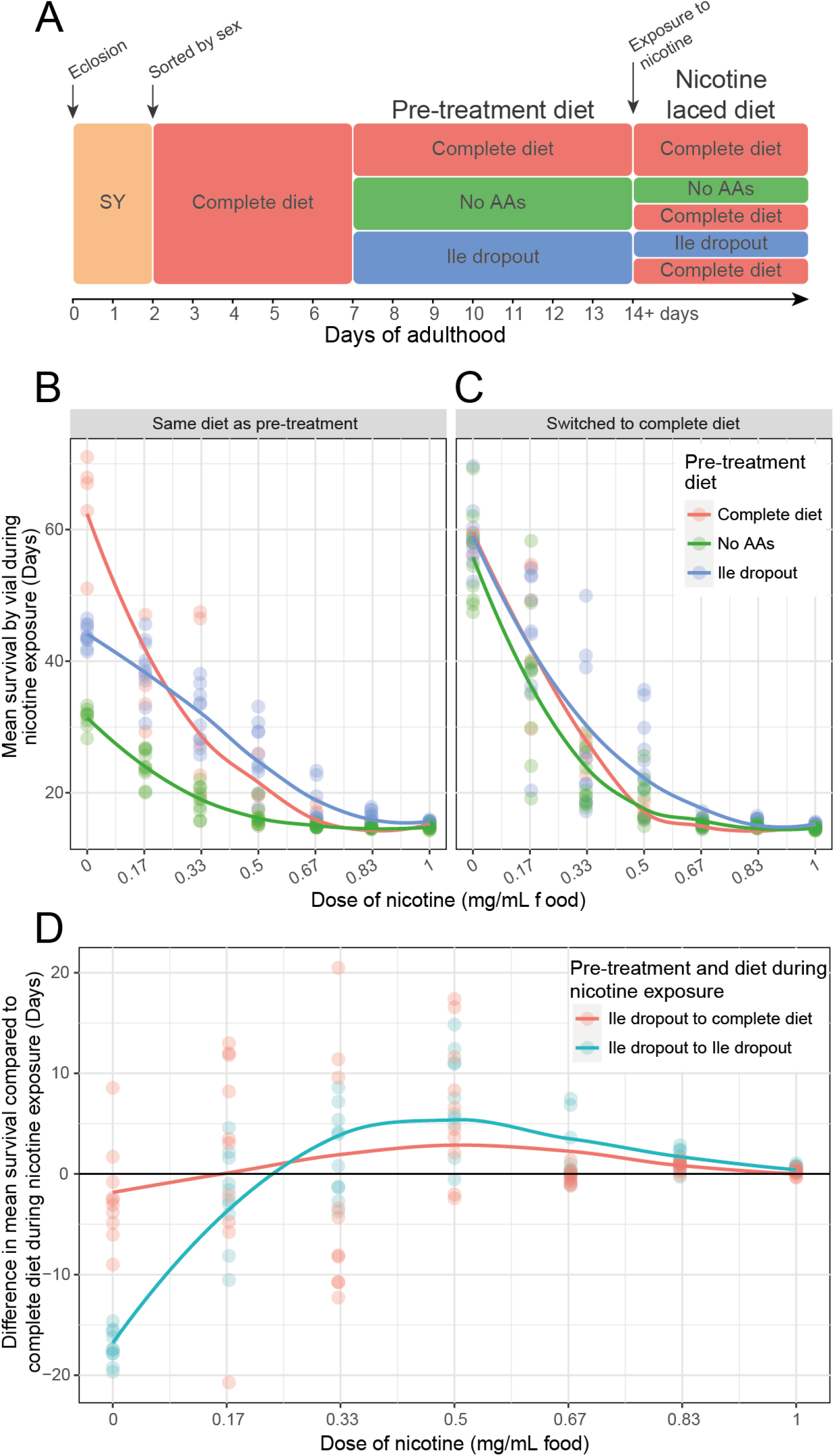
Pre-treatment modifies responses to higher doses of nicotine. (**A**) Flies were pre-treated with one of 3 diets (nutritionally complete, lacking all amino acids or lacking only isoleucine) for 7 days prior to chronic exposure of various nicotine concentrations. During nicotine exposure, flies were either (**B**) maintained on their pre-treatment diet or (**C**) switched to a nutritionally complete diet. (**D**) Difference between the mean survival of flies pre-treated with a complete diet and the survival of flies that were pre-treated with an isoleucine dropout and were either maintained on the same diet or switched to a complete diet at the time of nicotine treatment. N = 50 flies for each combination of diet and dose of nicotine.

Flies that were fed an isoleucine dropout diet showed resistance to doses of nicotine between 0.33mg/mL – 1mg/mL when compared to complete diet controls. When the nicotine concentration fell to 0.33 mg/mL, isoleucine deprivation did not extend life beyond that of the complete diet controls (∼30d mean, P = 0.75), and at lower doses (0.17mg/mL and 0 mg/mL) adult fly survival was compromised by dietary isoleucine omission, showing that the benefits of this diet for survival are only apparent when the flies are challenged with stronger nicotine stress.

Interestingly, although isoleucine is an essential amino acid, the flies on isoleucine dropout food always lived longer than those on diets lacking all amino acids (Figure 2.B), indicating again that eliminating single amino acids has different effects on the flies’ physiology than eliminating all amino acids. These data also indicate that any protective effects of isoleucine dropout are counteracted in the longer term by the negative effects of isoleucine omission. Thus, it is possible that fly survival may be further enhanced if flies were switched from the dropout pre-treatment diet to a complete diet at the point of nicotine exposure.

Switching flies from amino acid dropout food to complete food significantly improved the long-term survival of flies that were temporarily deprived of all amino acids or of isoleucine but not exposed to nicotine (Figure 2C; P < 0.001). However, for flies exposed to nicotine, there was no difference in protection when comparing flies that were maintained on their pre-treatment diet against those that were switched to a complete diet during the phase of nicotine exposure (Figure 2D; P = 0.57). These results suggest that it is the diet during pre-treatment that confers the benefit for survival during nicotine exposure. Once again, we observed no beneficial effect on nicotine resistance for flies exposed to diets lacking all amino acids, whether they were switched to a complete diet during nicotine treatment or not. Thus, the protective effect of isoleucine deprivation is confined to the pre-treatment period, during which time the flies require the presence of other dietary amino acids.

### Pre-treatment duration as well as amino acid identity and availability all modify protection against nicotine stress

Our previous data show that only specific single amino acid deprivations can enhance fly resistance to nicotine, and that protection is gained during the pre-treatment phase. To explore whether graded restriction of an amino acid was sufficient for protection, we restricted either threonine or isoleucine, to 75%, 50%, 25% or 0% of the amount in the complete diet. We opted to manipulate threonine in this assay as its removal previously showed a moderate protection (Figure 1C) and we wanted to explore whether this could be further increased. As we restricted the levels of these amino acids, we simultaneously investigated the relationship that pre-treatment duration had on protection by modifying the duration of dietary pre-treatment to 7, 5, 3 or 1 day prior to nicotine exposure (Figure 3A).

**Figure 3.**
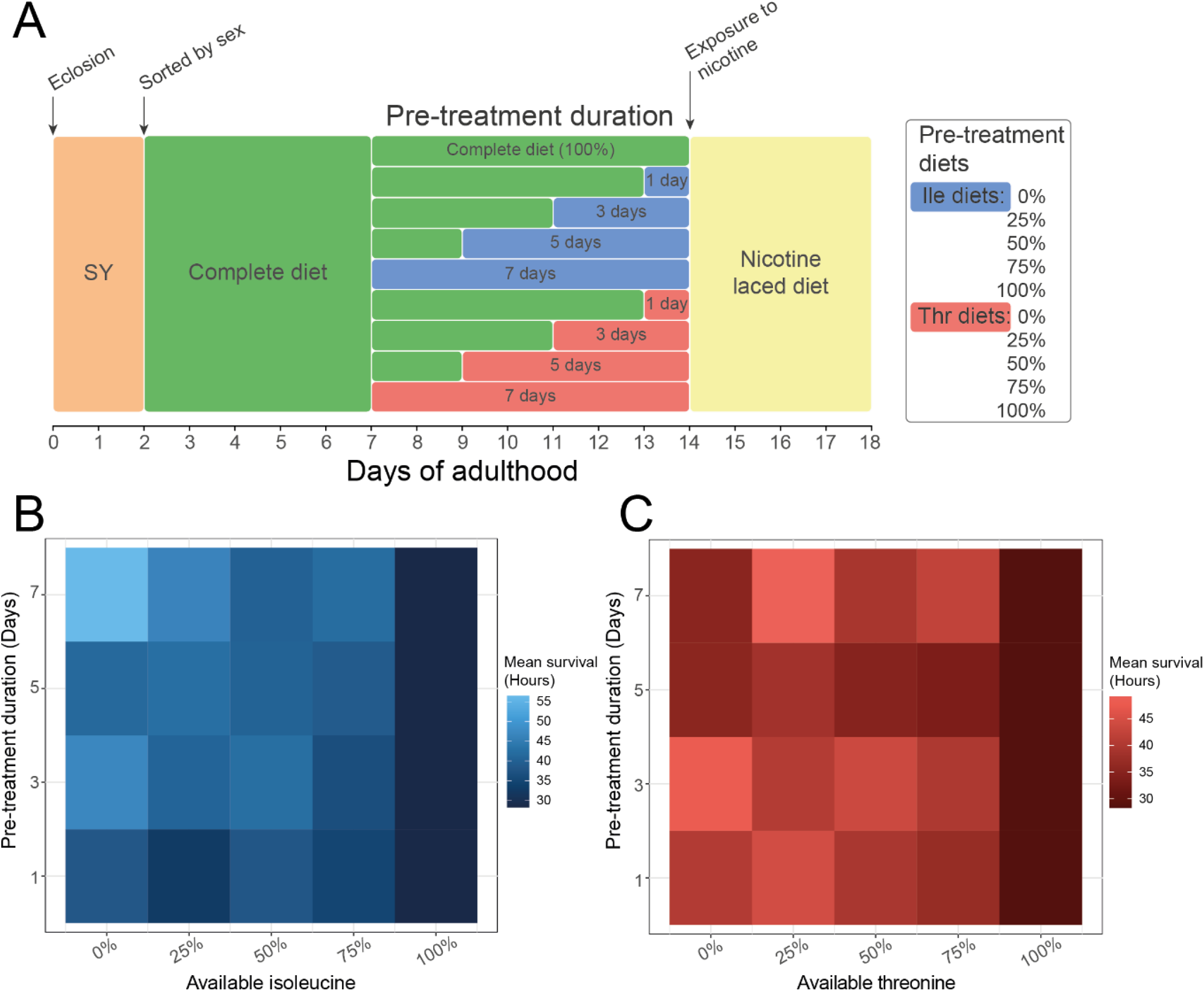
Pre-treatment duration as well as amino acid identity and availability all modify protection against nicotine stress. (**A**) Flies were pre-treated with diets containing 1 of 5 amounts of either isoleucine or threonine (0%, 25%, 50%, 75%, 100%) for either 7, 5, 3 or 1 day before chronic exposure to 0.83mg/mL nicotine. Mean survival of (**B**) isoleucine and (**C**) threonine modification depicted as heatmaps. N = 50 flies for each combination of pre-treatment duration and available amino acid. Pre-treatment with 100% of each amino acid (i.e. the complete diet) could not differ in duration time and N = 50 flies for this group in total.

Compared to the complete diet, we found that both the amount of available isoleucine and the pre-treatment duration modified protection during nicotine exposure (Figure 3B; P < 0.001). In general, survival improved in response to longer and more intense isoleucine dilutions such that the longest surviving flies were those pre-treated with 0% isoleucine for 7 days. However, when threonine was manipulated, we found that only the availability of amino acid (P<0.001) and not pre-treatment duration (P = 0.67) modified survival during nicotine exposure (Figure 3C). Interestingly, pre-treating flies with a diet containing 25% threonine was the most protective. Thus, the identity of the focal amino acid, the degree to which it is diluted, and, for some amino acids, the duration of exposure can all modify the way in which protection against nicotine is acquired. These data again show that while varying the concentration of different amino acids in the diet can elicit a similar phenotype, the dynamics of the protective effects are amino acid specific.

### Differences in survival during nicotine exposure between pre-treatment groups is not explained by whole body fat/protein composition or feeding behaviour

A potential explanation for the way amino acid manipulations protect flies against nicotine exposure is that a given pre-treatment diet could prime flies to simply consume less nicotine and therefore live longer. Using a Capillary Feeder (CaFe) assay (29) (Figure S2A), we found that the volume of pre-treatment medium consumed was indistinguishable between diets, except for increased consumption for flies on a diet lacking amino acids (Figure S2B). Importantly, compared to flies that were pre-treated with a complete diet, flies that were fed a diet lacking isoleucine ate as much nicotine laced medium (Figure S2C; P = 0.4) but were longer lived (P = 0.006). When analysing all diets together, we found that survival was modified by both the amount of nicotine consumed (Figure S2D; P < 0.001) and also by the type and amount of pre-treatment diet consumed (P = 0.03), but the relationship was positive, meaning that consuming more nicotine containing food correlated with surviving longer. This shows that it was beneficial for the flies to continue to eat food during nicotine exposure and that the protective effects of our pre-treatment diets were not because the flies simply ate less toxin.

Since our protective pre-treatment diets significantly reduced the egg production of flies without significantly reducing feeding behaviour, we anticipated that they would be under a positive energy balance and would therefore accumulate more body fat than controls. To assess if this were the case and if it contributed to nicotine resistance, we compared the amounts of triacylglycerol (TAG) and protein found in our flies on the same four dietary pre-treatments as we had used previously (Figure S3A). While flies on the amino acid-free or the isoleucine dropout diets had less whole-body protein than flies on the complete or the 25% threonine diet (Figure S3B; P < 0.03), neither whole-body fat content nor the ratio of fat:protein in these flies was significantly different across treatments (Figure S3C). These data indicate that body composition cannot explain the nicotine resistance phenotype.

### GCN2 and mTORC1 signalling confer protection against nicotine stress

Flies signal amino acid status via two evolutionarily conserved intracellular kinases: GCN2 and mTORC1. These act in a complementary fashion, with mTORC1 activated by the presence of amino acids, whereas GCN2 is activated by the absence of an amino acid. Thus, we would expect our pre-treatment diets to activate GCN2 and perhaps cause suppression of mTORC1. Although, to what extent mTORC1 is suppressed is not clear given that our diets still contained the amino acids that are known to be activators of mTORC1, including leucine, methionine, and arginine. To understand how changes in GCN2 and mTORC1 activity may be involved in protection against nicotine, we used genetic and pharmacological tools to suppress the activities of both kinases, individually and simultaneously, while also manipulating the diet composition (Figure 4A).

Overall, *GCN2* null flies were less resistant to nicotine than wildtype flies (Figure 4B; P < 0.001) and in contrast to wild type flies, neither isoleucine dropout nor threonine dilution induced protection against nicotine. By contrast, when wildtype flies were pre-treated with the mTORC1 suppressor rapamycin, nicotine resistance was elevated for all pre-treatment groups (Figure 4C; P < 0.02) except for those on the isoleucine dropout diet (P = 0.36), which were already protected when compared to flies on the complete diet (P < 0.001). Treatment with rapamycin also improved nicotine resistance in *GCN2* null flies that were pre-treated with either a complete diet or a 25% threonine diet (Figure 4D; P < 0.002), but not for *GCN2* null flies that were pre-treated with an amino acid-free diet or an isoleucine dropout diet (P > 0.2). Together these data indicate that intact GCN2 is required for dietary protection against nicotine stress and that mTORC1 suppression is sufficient for protection. They also indicate that the isoleucine dropout may act via mTORC1 suppression, but only when GCN2 activity is intact. By contrast, the more subtle protection offered by threonine dilution can be augmented by mTORC1 suppression, whether or not GCN2 is intact. These data point to a complex interaction whereby dilution of individual amino acids confer protection against nicotine by tuning GCN2 activation and mTORC1 suppression.

**Figure 6.**
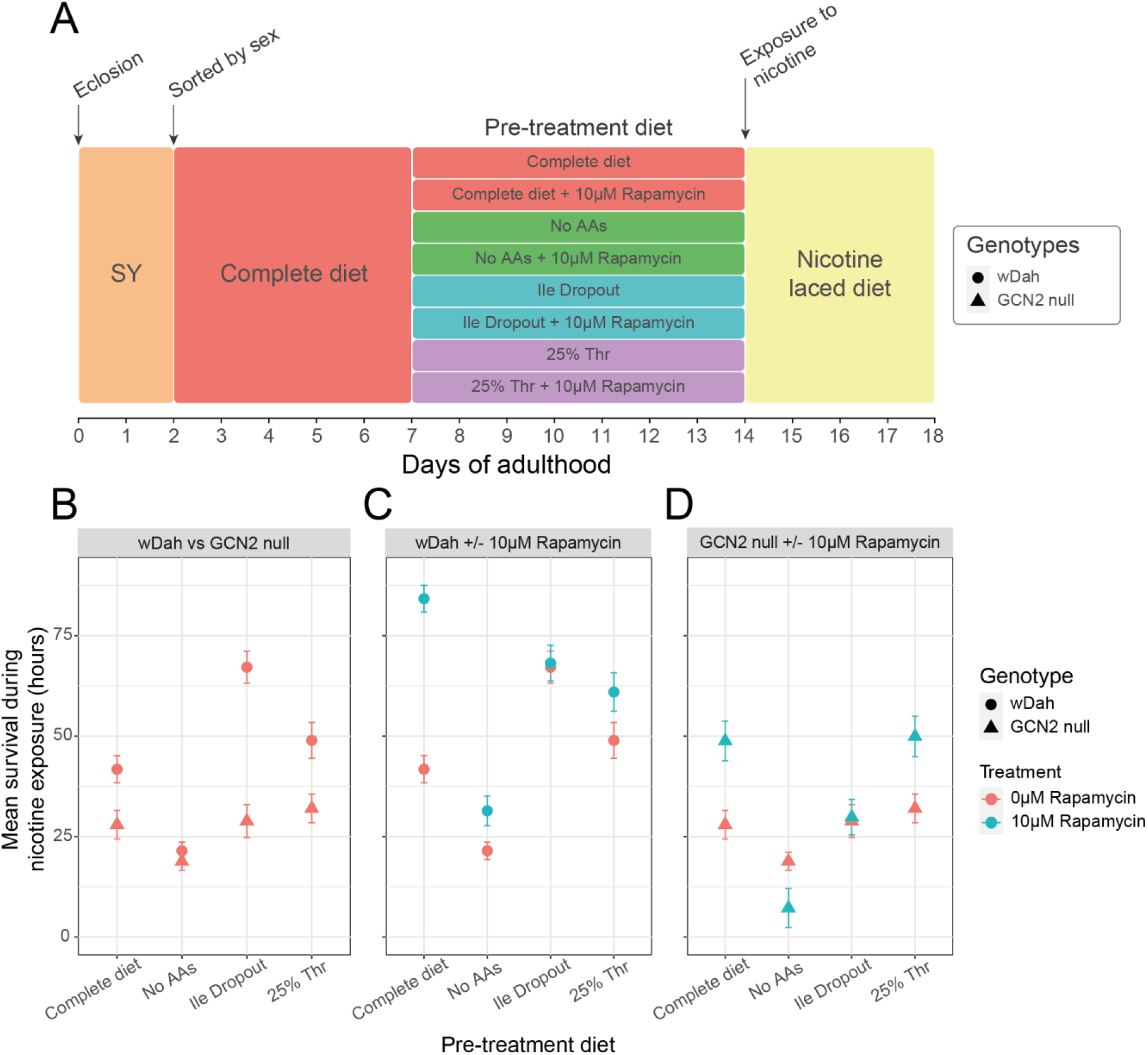
GCN2 is necessary for dietary protection against future nicotine stress and suppression of mTOR is sufficient to induce protection. (**A**) Prior to chronic exposure to 0.83mg/mL nicotine, GCN2 null and wDah control flies were pre-treated for 7 days with 1 of 4 diets that either contained 0µM or 10µM of the mTOR suppressor rapamycin. Mean survival +/- SEM of (**B**) wDah and GCN2 nulls, (**C**) wDah pre-treated with food containing either rapamycin- or vehicle (ethanol), and (**D**) GCN2 nulls pre-treated with rapamycin- or ethanol-laced food. N = 50 flies for each combination of genotype, rapamycin concentration and pre-treatment diet.

## Discussion

Treating organisms with nutritionally modified diets has been shown to increase stress tolerance in different model systems (10-12), although the exact mechanisms by which this occurs remain elusive. In this paper, we expand these findings by establishing that briefly exposing *Drosophila* to amino acid manipulated diets can enhance their tolerance of a chemical stressor. This type of protection is an example of cross-modal hormesis whereby a mild dietary stress can elicit protection against another, apparently unrelated, stressor (30). Importantly, we found that the magnitude of protection depends on several factors, including the identity of the manipulated amino acid, the intensity of deprivation, and the duration of pre-treatment. We also found that this protection is mediated by interactions between the nutrient sensing kinases GCN2 and mTORC1. These data provide insights into the way diet and nutrient signalling can be harnessed to manipulate an evolutionarily conserved aspect of stress protection. This has important implications for pest control in agriculture as well as for animal and human health.

Amino acids are both building blocks for protein biosynthesis and required to trigger signalling to regulate proteostasis and anabolism so that physiology is tuned to match the supply of available nutrients (31). Changes in the amount and relative abundance of dietary amino acids can modify organismal behaviour, growth, reproduction, and lifespan, but which amino acids are key, and for what reason, is not known. One of the more well studied amino acid manipulations is methionine restriction, which can enhance stress resistance and lifespan in rodents (11, 12, 32, 33), and can extend lifespan in flies (34, 35). But methionine restriction may not be special in this regard since further work in flies has since shown that lifespan can also be enhanced by simultaneously restricting the three branched chain amino acids (BCAA; isoleucine, leucine, and valine) or by restricting another set of three essential amino acids (threonine, histidine, and lysine) (36). Together, these data show that manipulations of various dietary amino acids can modify the consumer’s physiology and fitness, but because we do not fully understand the biological roles of each amino acid, we cannot deduce the logic by which each may be important for a particular phenotype. In our study, transiently depriving flies of each one of the essential amino acids revealed a protective effect against nicotine that ranged from ∼40-100% longer life following isoleucine deprivation to no protection at all following methionine or leucine deprivation. It will be interesting in future studies to determine the breadth of this protection as additional phenotypes may provide useful in understanding the metabolic physiology of amino acids.

Of the biochemically related groups of amino acids in our experiments, we thought it particularly interesting that deprivation for each one of the three BCAA elicited a different protective response that ranged from the strongest protection (isoleucine) to intermediate protection (valine) to no protection (leucine). The BCAAs are often studied together as a group because of their role in anabolism, health, and signalling (37), and there has been a lot of attention on how dietary BCAA levels alter metabolic health and longevity in mammals (38). Interestingly, Yu, Richardson (39) recently showed that restriction of isoleucine or valine, but not leucine, can improve metabolic health in both normal and obese mice. These findings directly parallel the protective effects against nicotine of each of the BCAA restrictions in our flies, indicating a possible mechanistic overlap. However, we also note that in another recent study, Yap, Rusu (40) found that protection against obesity-related defects in mice was conferred by restriction of dietary threonine or tryptophan, but not the BCAAs. Perhaps small differences in treatment conditions, such as duration, intensity, and dietary context, have implicated different amino acids as beneficial in these two mouse studies in the same way that we have observed their effects can vary for flies. As we home in on the mechanisms by which our amino acid diets afford protection against nicotine, we will gain insights into whether these same processes can also protect against the metabolic dysfunction associated with obesity.

The two major evolutionarily conserved amino acid sensors, mTORC1 and GCN2, initiate signalling in response to opposing nutritional signals. mTORC1 is activated by amino acids, while GCN2 becomes active in the absence of amino acids (41). These complementary signals work together to maximise the use of amino acids for anabolism when they are available, and to protect cells against deficiencies for these essential nutrients. We found that pharmacological suppression of mTORC1 on a nutritionally complete diet could phenocopy the maximum protective effects of transient amino acid deprivation. We also found that when flies were null for GCN2, neither isoleucine deprivation nor threonine restriction were protective. Further, the beneficial effects of rapamycin were dampened in GCN2 null flies. These results indicate that both kinases contribute to protect flies against nicotine, and that functional GCN2 is required to promote the full protective effects of mTORC1 suppression.

Past work with bees (*Apis mellifora*) and the tobacco hornworm (*Manduca sexta*) has shown that nicotine detoxification is achieved via enzymes in the drug detoxification system that act to detoxify and excrete xenobiotic compounds (42, 43). Many of the genes important for this system are under the control of the Nrf2 (nuclear factor erythroid 2–related factor 2) transcriptional regulator, which binds to antioxidant response elements (AREs) in the regulatory regions of these genes (44, 45). Our data indicate that reduced mTORC1 and activated GCN2 may work together to activate this system, potentially by activating Nrf2 (46). If this is the case, we expect that transient deprivation for any one of the amino acids that induced nicotine protection would also induce protection against other xenobiotics that are neutralised by this system. In fact, activating the drug detoxification pathways may also protect consumers against detrimental endobiotic toxins that arise as a by-product of metabolism. If true, this type of system could explain how low dietary amino acids can protect against acute exposure to a toxin as well as provide benefits to long term health in the absence of toxins (47).

An interesting new finding in our study is that both dietary concentration and duration of exposure determine the efficacy for protecting the consumer against nicotine. One explanation for this is that these conditions differentially modify signalling activity of mTORC1 and GCN2, and this results in different levels of protection. This is a plausible explanation as we know that different amino acids change both mTORC1 activity and GCN2 activity to different extents (41, 48). We also note that while signalling triggered by amino acid dilution may be beneficial, the absence of an essential amino acid may have an antagonistic effect on consumer physiology and so limit protection. This could vary by amino acid and would be a function of the extent to which each amino acid is required for protection, the amount of the amino acid that can be safely scavenged from body stores, and the duration over which protective amino acid repurposing is required. Evidence for this can be found in our data: 7 day, short term, deprivations of isoleucine protected flies against nicotine, but longer-term deprivation was detrimental to health and shortened lifespan, even in the absence of nicotine. A similar situation could be envisaged for the lack of protection we observed after methionine deprivation in our study, which contrasts its protective effects when restricted in other studies (11, 32, 49). Methionine is both required for the synthesis of new proteins as well as being a source of cysteine to produce glutathione. Glutathione is an important peptide for conjugation to xenobiotics in phase II detoxification and its synthesis is upregulated in bees exposed to nicotine (42). Thus, removing methionine from the diet may stimulate the signals for protection, but at the same time prevent the production of peptides and proteins that are required for resistance. A full characterisation of this phenotype will therefore not only require a better understanding of the signalling involved, but also quantification of the amino acid requirements to build the protective systems being activated.

## Conclusion

Our data reveals new insights into the way that dietary amino acid balance modifies organismal health. We found that restricting flies of an essential amino acid can protect them against subsequent chemical stress in a way that depends on interactions between the identity of the amino acid, the intensity of deprivation, and the duration of pre-treatment. This protection is mediated by interactions between GCN2 and mTORC1. This cross-modal hormesis paves the way for a better understanding of the environmental factors that alter insect susceptibility to pesticides and could open opportunities for the use of diets to help patients recover from planned treatments with toxins such as chemotherapy.

## Methods

### Fly husbandry

Experiments were conducted with adult, female, mated white-eyed *Drosophila melanogaster* (strain Dahomey; wDah) unless otherwise specified. wDah stocks are maintained outbred in continuous overlapping generations housed in a high-density population cage at 25°C, ambient humidity and a 12:12 hour light:dark cycle. The *GCN2* null *Drosophila melanogaster* used were a gift from Dr. Sebastian Grönke and Professor Linda Partridge (Max Planck Institute for Biology of Ageing) and we introgressed the *GCN2* null transgene into our wDah background. The GCN2 null stock is maintained at 18°C, 60% humidity and a 12:12 hour light:dark cycle. All stocks are maintained on a sugar yeast (SY) diet (50).

### Synthetic media

Chemically defined synthetic (holidic) media used in the experiments were prepared as described in Piper, Blanc (23) using the exome matched (FLYaa) ratios of amino acids as described in Piper, Soultoukis (51). Holidic media were prepared in advance and stored for up to 4 weeks at 4°C until required.

Nicotine-laced media were prepared by aliquoting 100µL of free-base nicotine [Sigma-Aldrich: N3876] diluted in absolute ethanol onto the surface of cooled holidic media. The media were then kept at room temperature for 2 days to allow for the dispersion of nicotine before immediate use. Freshly-laced media were prepared as required and were not stored.

Rapamycin-laced media were prepared by aliquoting 30µL rapamycin [Jomar: S1039] dissolved in absolute ethanol onto the surface of cooled holidic media. The media were then kept at room temperature for 1 day to allow for the dispersion of rapamycin before storing at 4°C for up to 7 days until required.

Liquid holidic media used in CaFe assays were prepared by omitting agar and they were stored in falcon tubes at −20°C until required. Nicotine-laced liquid holidic medium was prepared by aliquoting 9.96µL of 100% free-base nicotine into 12mL liquid medium (0.83mg/mL).

### Nicotine assays

Age-matched cohorts of flies were generated for experiments in controlled population density using the methods described by Linford, Bilgir (52). Briefly, adult flies were kept on apple juice agar plates with live yeast paste and allowed to lay for 24h, after which the eggs were collected in PBS and pipetted onto SY medium. Experimental flies were reared from egg to adult using SY medium and transferred to fresh SY medium as a mixed sex group to mate for 48 hours post eclosion. Flies were mildly anaesthetised with CO_2_ on day 2 of adulthood to be sorted by sex onto a complete synthetic diet. Unless otherwise specified, flies were transferred to their pre-treatment diet on day 7 of adulthood and transferred to nicotine-laced medium on day 14 of adulthood. During pre-treatment, survival was recorded using the software DLife (52) as flies were transferred to fresh vials each Monday, Wednesday, and Friday. When exposed to nicotine, survival was recorded 3 times a day for 5 days or until all flies were dead. Experiments were conducted in controlled conditions: 25°C, 70% humidity, and a 12:12 hour light:dark cycle.

### Capillary Feeder (CaFe) assay

Flies for the CaFe assay were generated in the same way as for nicotine assays. On day 7 of adulthood, flies were individually aspirated into vials containing 3mL of agar and a capillary [Sigma-Aldrich: Z543241] containing 5µL of their assigned pre-treatment diet. Consumption was measured for each fly every 24h, and the capillaries were blotted, rinsed, and refilled with fresh medium at this time. On day 14 of adulthood, all flies were presented with capillaries containing a complete medium laced with 0.83mg/mL nicotine. Food consumption continued to be measured every 24h during treatment with nicotine and survival was recorded 3 time a day. To control for evaporation, volume lost from capillaries kept in vials with no flies was also measured. CaFe assays were conducted in controlled conditions: 25°C, 80% humidity and a 12:12 hour light:dark cycle.

### Whole-body triglyceride and protein content assays

Flies for the body composition assays were generated in the same way as for nicotine assays. On day 14 of adulthood, after 7 days of pre-treating with diet, groups of 5 flies were aspirated into cryovials, frozen in liquid nitrogen, and stored at −80°C until measured. Frozen flies were transferred to 1.5mL Eppendorf tubes with 500µL 0.05% Tween 20 in water. Using a pestle, flies were homogenised then an additional 500µL of 0.05% Tween 20 was added and the samples vortexed for 20 seconds. Samples were incubated at 70°C for 5 minutes and then centrifuged at 5000rpm for 1 minute. The supernatant was collected and centrifuged again at 14,000rpm for 3 minutes. Finally, the supernatant was again collected and used for measurements.

For triacylglycerol (TAG) measurements, 50µL of sample or a dilution series of a glycerol standard was aliquoted into a 96 well plate. Samples were aliquoted in triplicate for TAG measurements and an additional well was aliquoted as a sample blank. 200µL of Infinity™ Triglycerides liquid stable reagent [ThermoFisher: TR22421] was added to each well containing sample, except blanks, to which 200µL of water was added.

For protein measurements, 10µL of sample or BSA standard was aliquoted in triplicate into the same 96 well plate as TAG samples. The Pierce™ BCA Protein Assay Kit [ThermoFisher: 23225] was used according to the manufacturer’s instructions. Samples were measured for absorbance at 540nm for TAG and 562nm for protein.

To calculate the whole-body composition of flies, the absorbance values of TAG blanks were subtracted from the average of each TAG triplicate. The mass of TAG per fly was then calculated from the standard curve and then normalised by dividing by the average mass of protein per fly.

## Figures and Data Analysis

We used R version 4.1.1 (53) to conduct all analyses and we created all plots using ggplot2 (54). All data and scripts are publicly available at [To be made freely available through Figshare].

Survival analysis was completed using Cox Proportional-Hazards modelling. We used the “coxph” function from the package survival (55) to model diet as a function of time to death after initial exposure to nicotine.

To measure differences in fecundity, and also whole-body fat and protein content, we fit a regression with pre-treatment diet as the independent variable in all cases. We then modelled these data using the base R function “aov”. Significantly different pairs of treatments were determined by performing a post-hoc Tukey’s HSD test on these models using the base R “TukeyHSD” function.

Nicotine dose response curves were modelled using a self-starting exponential model from the package drLumi (56). A limitation of this type of modelling is that only 2 models can be compared at once. We generated comparisons in a fully factorial manner and then adjusted P values to account for multiple tests using a Bonferroni correction.

Linear models (base R, “lm”) were used to model the effect of diet on the differences in mean survival across nicotine doses. We first normalised the data relative to the complete diet, fixing the response of the complete diet to y = 0. We then compared how switching or not switching diet during nicotine exposure affected normalised survival across nicotine doses.

To find parsimonious models that best fit our data, we compared two nested models at a time using base R’s “anova”. These models were nested such that only one variable was removed from the more complex model to form the reduced model. If the models were not significantly different from each other, the model with the lowest “AIC” (base R) value, typically the reduced model, was used for further analysis. We found that a second order polynomial linear model best fit these data, and the diets were compared as a factor within the same model using “emtrends” from the emmeans package (57). We then determined whether these trends were significantly different from each other using “cld” from the multcomp package (58). To determine if these diet switch models were different from the complete diet, we used “emmeans” from the emmeans package (57) and tested if the models were different from 0.

Linear models (base R, “lm”) were also used to model both duration of pre-treatment and dose of amino acid as a function of mean survival. When analysing these relationships, we found that second order polynomial models best fit the data using the process outlined above. To determine whether pre-treatment duration or availability of the focal amino acid significantly impacted survival, we analysed the model using a type II ANOVA from the package car (“Anova”; Fox and Weisberg (59)).

## Acknowledgements

We thank Dr. Sebastian Grönke and Professor Linda Partridge (Max Planck Institute for Biology of Ageing) for allowing us to use unpublished reagents, the *GCN2* null fly line, in our experiments and this manuscript. Additionally, we extend thanks to Amy Dedman, Brooke Zanco and Krithika Balagopal (Monash University) for assistance with data collection during the peak of Melbourne’s COVID19 lockdown restrictions. We are also grateful to Joshua Johnstone (Monash University) for help with backcrossing lines and maintaining stocks. Finally, we would like to thank Andrea Chan (Monash University) for her involvement in the initial stages of this project.

## Supplementary figures

**Figure S1.**
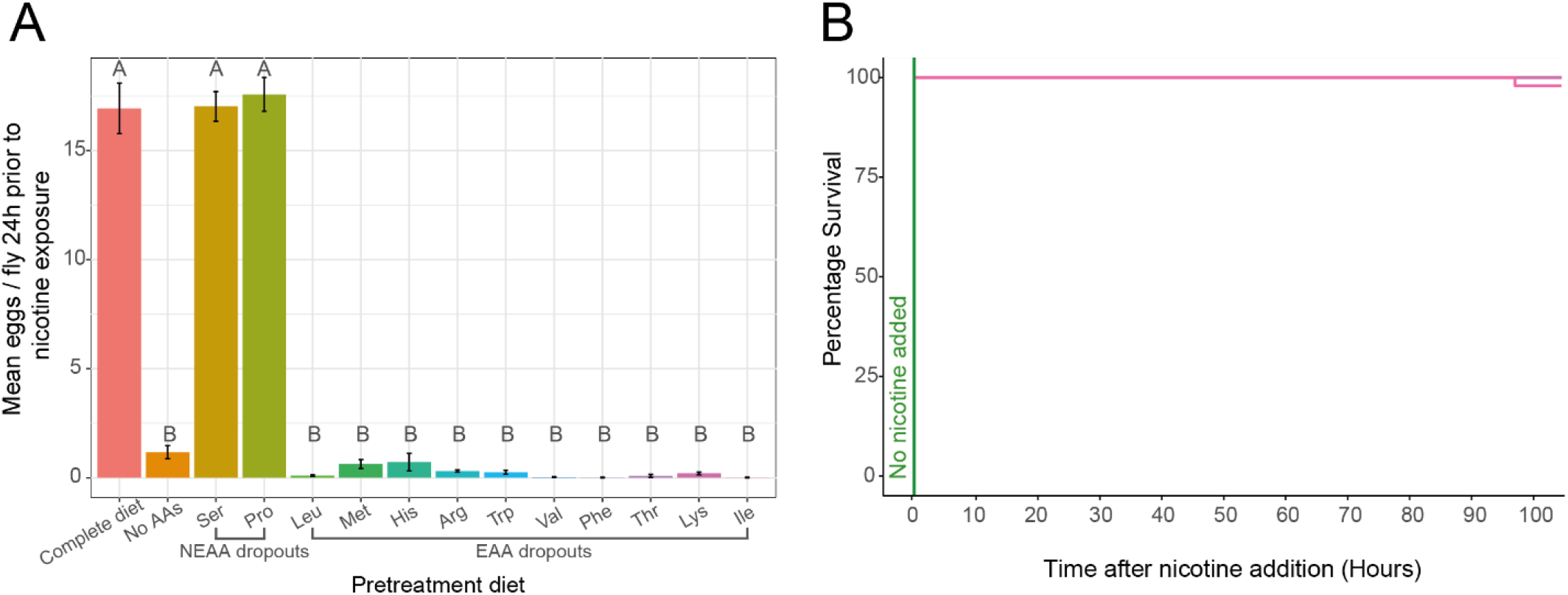
Responses to removing non-essential and essential amino acids from the diet. (**A**) Fecundity prior to nicotine exposure did not statistically differ between the non-essential amino acid dropouts and the complete diet (P > 0.98). Removing any one of the essential amino acids, or when removing all amino acids, there was a near complete suppression of egg laying that was no different between the different dropout diets (P > 0.50; comparing amongst single essential amino acid dropouts and diet containing no amino acids). (**B**) Flies that were pre-treated with a dropout diet and not exposed to nicotine did not suffer deaths during the assay period. N = 50 flies that were pre-treated with a complete diet and N = 100 for all other pre-treatment groups.

**Figure S2.**
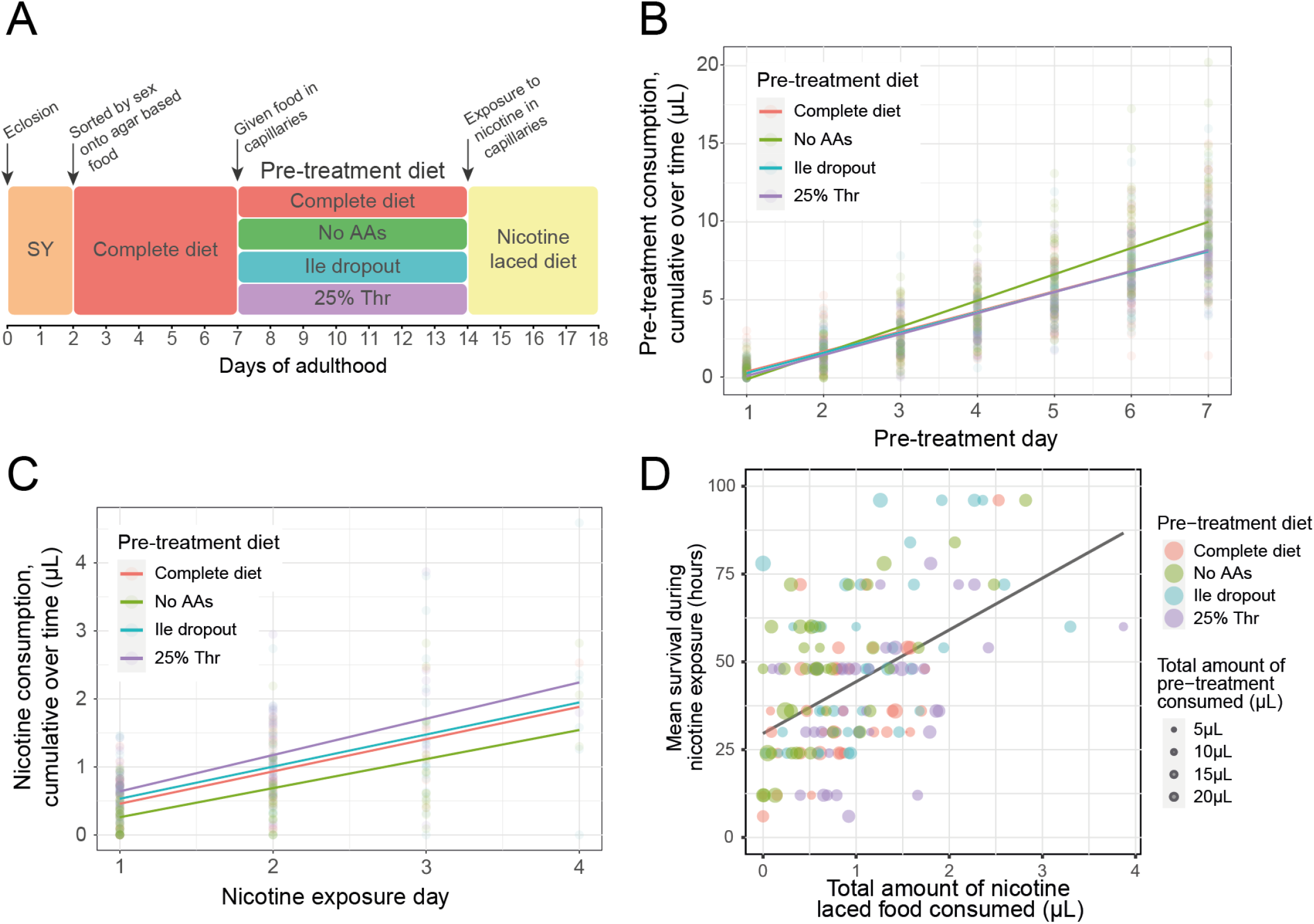
Improved survival during nicotine exposure was not caused by the flies eating less nicotine-laced food. (**A**) Flies were individually housed and pre-treated with 1 of 4 diets for 7 days prior to chronic exposure to 0.83mg/mL nicotine. All food was delivered via capillaries and consumption was measured every 24h. (**B**) Flies that were pre-treated with an amino acid-free diet ate significantly more food than flies that were pre-treated with other diets (**C**) Compared to flies pre-treated with a complete diet, flies that were pre-treated with a 25% threonine diet ate significantly more nicotine laced medium, whereas flies pre-treated with an amino acid-free diet consumed less and those pre-treated with the isoleucine dropout diet ate the same amount. (**D**) Survival was modified by the total amount of nicotine consumed and this was modified by the type and the amount of the pre-treatment diets consumed. N = 40 flies per pre-treatment group.

**Figure S3.**
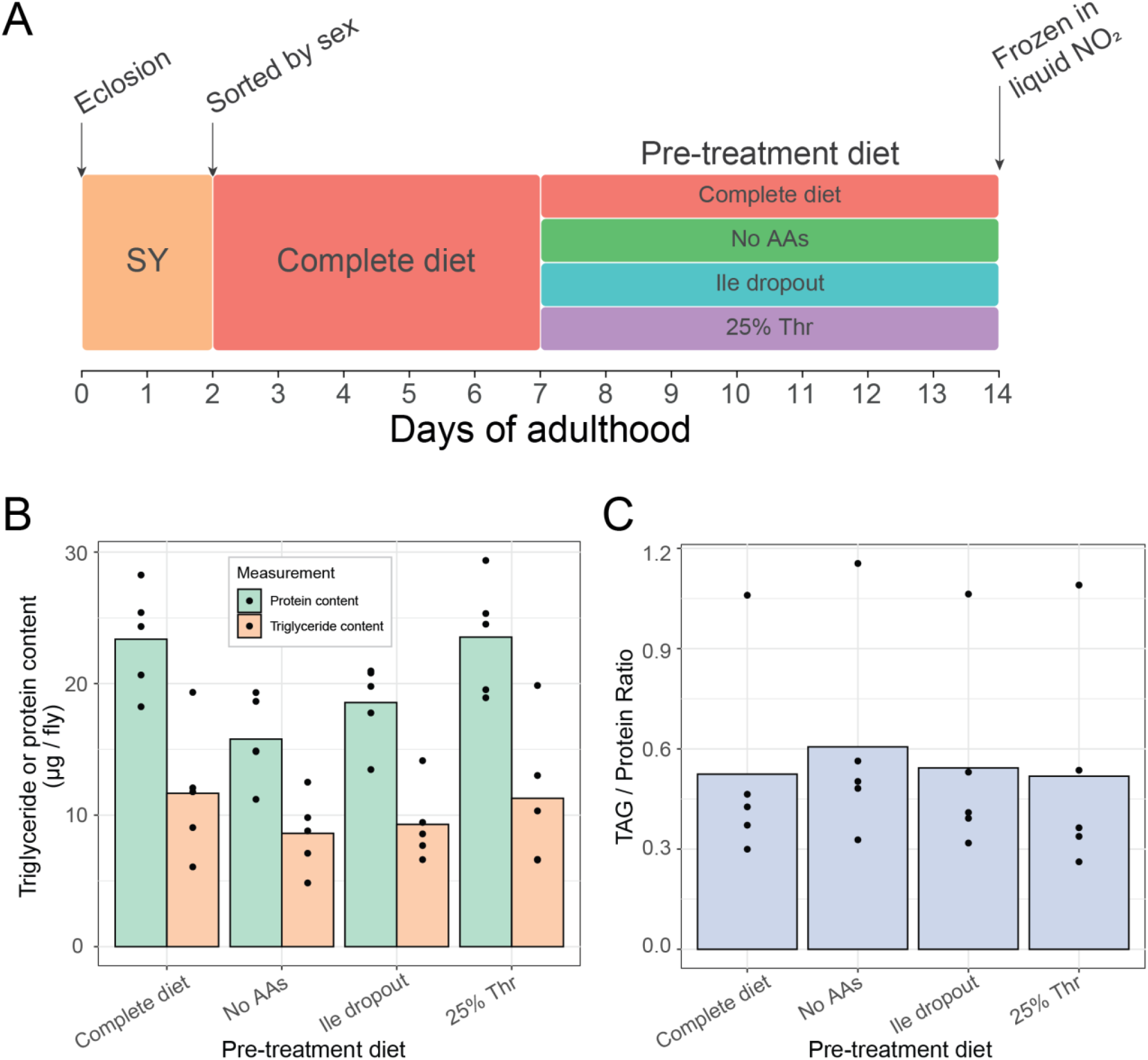
Differences in survival during nicotine exposure between pre-treatment groups is not explained by whole body fat or protein contents. (**A**) Flies were pre-treated with one of 4 diets for 7 days prior to being frozen in liquid nitrogen for whole-body protein and fat assays. (**B**) Flies that were pre-treated with an amino acid-free, or an isoleucine dropout diet had significantly less protein than flies pre-treated with either a complete diet or a diet containing 25% threonine (P < 0.03). Pre-treatment diet did not significantly impact whole-body fat content (P > 0.1). (**C**) When whole-body fat content was normalised to protein content, no significant differences were found between the flies from different pre-treatment diets (P > 0.8). N = 5 per pre-treatment group where each replicate is the mean value from 5 individual flies.

